# Conjugation Based CRISPR Antifungals (COBRA)

**DOI:** 10.64898/2025.12.06.692781

**Authors:** Vida Nasrollai, Bogumil J. Karas

**Affiliations:** Department of Biochemistry, Schulich School of Medicine and Dentistry, Western University, London, ON N6A 5C1, Canada

**Keywords:** Conjugation, CRISPR, Cas9, antifungal, *S. cerevisiae*, COBRA

## Abstract

Fungal infections are increasingly difficult to treat due to rising antifungal resistance and the limited number of effective drug classes. To address this challenge, we developed a conjugation-based CRISPR antifungal (COBRA) platform that enables delivery of programmable gene-targeting machinery from *Escherichia coli* to *Saccharomyces cerevisiae*. We engineered a mobilizable pVenom plasmids containing an oriT for conjugation, Cas9 and guide RNAs for elimination of yeast. To prevent toxicity in *E. coli*, yeast *ACT1* intron is inserted in *Cas9*. Seven guide RNAs targeting essential genes involved in cell cycle progression, ribosome function, and DNA replication were first assessed by using electroporation as delivery and demonstrated strong lethality for guides targeting *CDC28* and *MCM2*. When delivered by conjugation, these guides again reproduced these results, whereas an intron-targeting control remained non-lethal. Dual-guide plasmids eliminated yeast colony formation entirely. Together, these findings demonstrate that conjugation-delivered CRISPR machinery can successfully target endogenous fungal genes and, especially when multiplexed, offers an effective and programmable antifungal strategy.

**Graphical Abstract:** 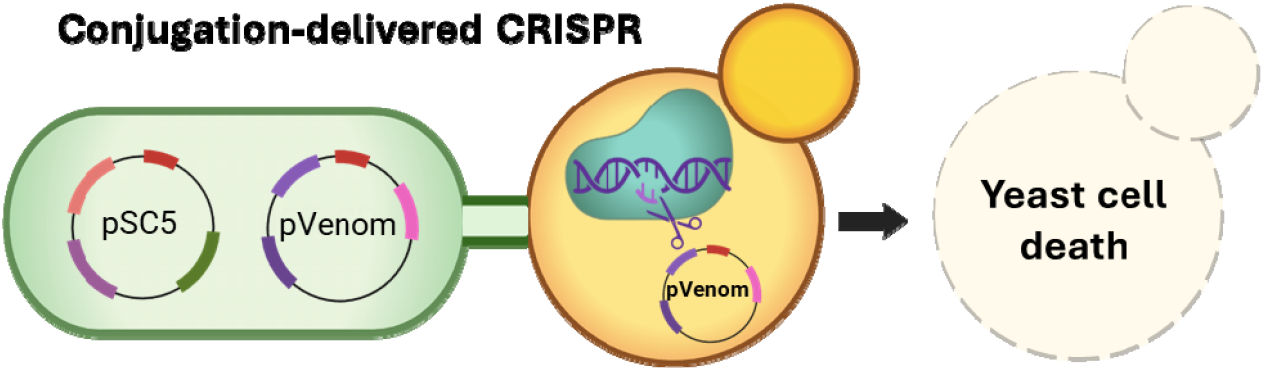

## INTRODUCTION

Fungal pathogens have become an important global health concern, particularly for people with weaker immune systems. Systemic infections have high mortality,^1^ and treating invasive fungal disease is still difficult because diagnostic tools are limited and current therapies are usually not enough to inhibit them.^2,3^ As antifungal resistance increases, it is even harder to cope with this problem, resulting in longer treatments and extended hospital stays.^4,5^ Current antifungal drugs fall into four main classes: azoles, pyrimidines, polyenes and echinocandins.^6,7,8^ However, resistant strains have emerged against all of these classes. For example, *Candida auris* has gained significant attention over the past decade as an emerging fungal pathogen with multidrug resistance to available antifungal agents^1^. Its rapid and widespread emergence has made it a global public health concern. Various strategies have been explored to address rising antifungal resistance, including large-scale screening efforts to identify new antifungal compounds.^9^ Although these methods were useful in earlier drug discovery, recent screens commonly identify known chemical scaffolds, showing how difficult it is to uncover new antifungal agents.^10^ Consequently, there is an urgent need for development of alternative strategies to combat fungal resistance and expand the therapeutic toolkit.^11,10^ Current antifungal therapies focus on a narrow set of conserved cellular targets, which restricts their specificity and makes resistance easier to develop.^7^. These agents can also cause off-target toxicity due to similarities between fungal and human cells.^2^

Bacterial conjugation enables the direct transfer of DNA between cells and can occur between different species, allowing plasmids to move into diverse bacterial and eukaryotic hosts.^12^ Although originally it has been thought that conjugation is restricted to prokaryotes, work by Heinemann and Sprague (1989) demonstrated that *E. coli* can transfer mobilizable plasmids into *S. cerevisiae* by conjugation, showing the possibility of inter-kingdom DNA exchange.^13^ Since then, conjugation has been shown to function across even broader eukaryotic hosts, including algae, and multiple fungal species^14,15,16,17,18,19,20,21,22^. Recent studies have also showed that conjugative *E. coli* donors can efficiently transfer mobile plasmids to a broad range of microbial recipients, including both Gram-negative and Gram-positive bacteria.^23^ Conjugation-based transfer into yeasts and pathogenic fungi has also been demonstrated, with *E. coli* conjugative plasmids into *Candida, Metschnikowia*, and multiple *Saccharomyces* strains^21^, as well as the filamentous plant pathogen *Ustilago maydis*.^*22*^ These properties allow conjugation to be used as a programmable method for introducing engineered DNA into fungal recipients. Conjugation-based antifungal activity has been previously demonstrated. Cochrane *et al*. (2022) showed that pSC5-based conjugative plasmids can deliver genetic elements, including toxic modules, into *S. cerevisiae*, indicating that conjugation can function as a useful approach for killing target fungi.^21^ More recently, Stindt and McClean^24^ showed that interdomain conjugation can deliver a CRISPR machinery that depresses or even collapses yeast growth, it is supporting the idea that conjugation can be used to eliminate fungal recipients. Additionally, conjugative plasmids have been shown to disseminate programmable CRISPR systems across diverse microbial species, highlighting their versatility as mobile genetic platforms.^23,25^ CRISPR nucleases introduce targeted double-strand breaks at defined genomic loci, and in *S. cerevisiae*, a Cas9-induced break at a specific site produces detectable toxicity and reduces cell viability.^26^ Because Cas9 generates a site-specific double-strand break, successful genome editing requires repair of this chromosomal damage using a supplied donor DNA.^27^ Here, we extend these concepts by developing a conjugation-based CRISPR antifungal (COBRA) technology that directly targets endogenous fungal genes. We engineered and validated mobilizable plasmids carrying CRISPR machinery that can be efficiently transferred from *E. coli* donors into *S. cerevisiae*, where they act as a potent, programmable antifungal reagent. This work establishes a foundation for next-generation targeted antifungal strategies, particularly those leveraging multiplexed gene targeting.

## RESULTS & DISCUSSION

To test whether conjugation-delivered CRISPR machinery could kill *S. cerevisiae* cells by targeting endogenous genes, we constructed a mobilizable pVenom (pV) plasmids (Figure 1A). These plasmids carry Cas9 nuclease with a yeast *ACT1* intron^28^ to prevent Cas9-associated toxicity in *E. coli*. It also carries an origin of transfer (oriT) required for mobilization from a bacterial donor into *S. cerevisiae* and a guide RNA expression cassette targeting specific genes (pV-g1 −7) or control plasmid (pV-GFP). In our design all pVenom plasmids are mobilized using the conjugative plasmid pSC5^21^ (Figure 1A). To assess whether our system could function as an antifungal, we established a donor system in which *E. coli* cells carry both the pSC5 (gentamicin selection in *E. coli*) and pVenom (Kanamycin selection in *E. coli*) plasmids. In yeast pSC5 (which is self-transmissible) can be selected on media lacking histidine or supplemented with nourseothricin or both, and either pV-GFP or pV-g1-7 plasmids can be selected on media lacking uracil (Figure 1B).

**Figure 1.**
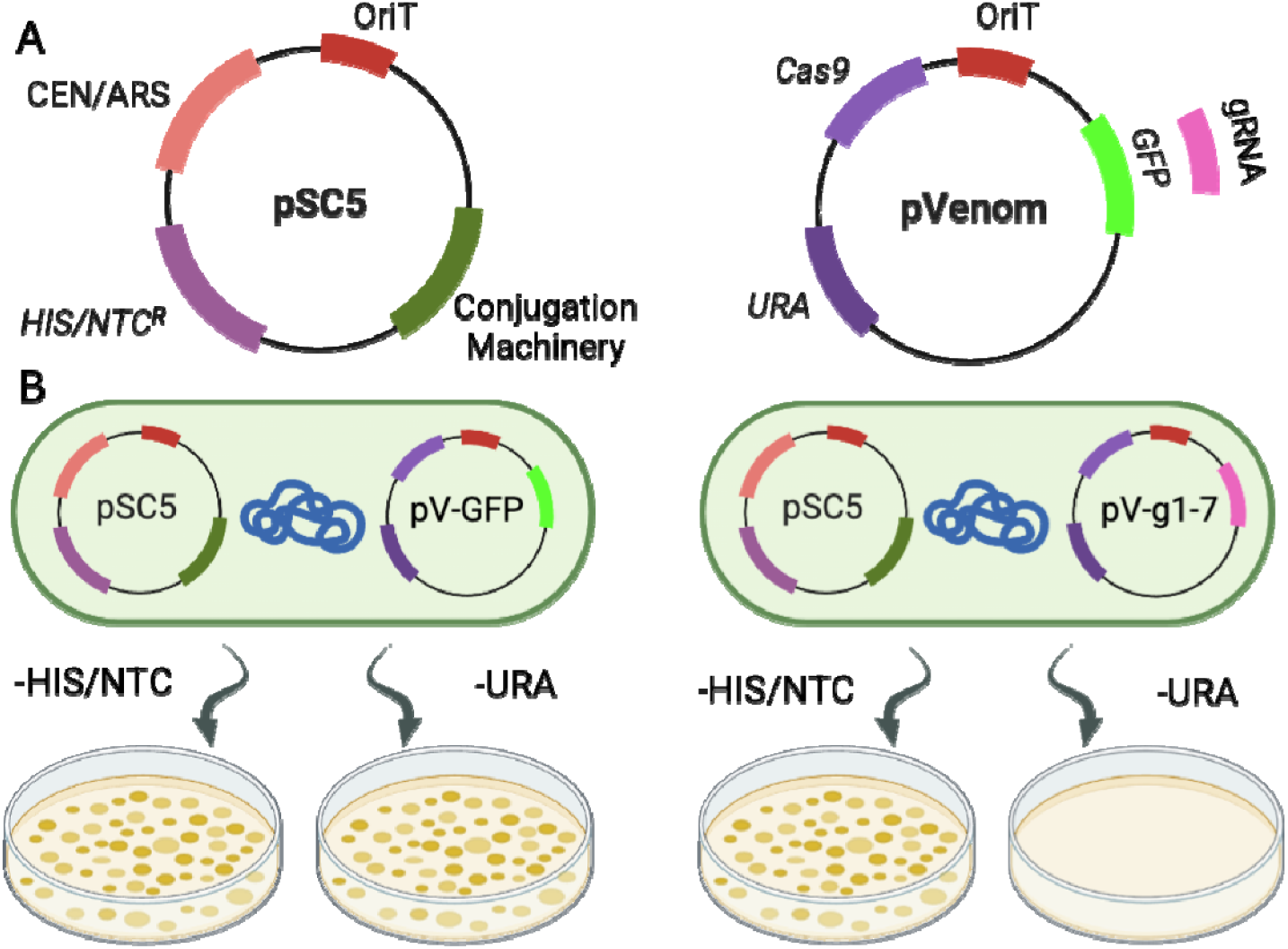
Conjugation-based CRISPR antifungal technology. (A) Plasmid maps of pSC5, pVenom plasmids. OriT = origin of transfer, CEN = yeast centromere, ARS = autonomous replication sequence, HIS = histidine selection marker, NTC = nourseothricin resistance gene, Cas9 = CRISPR-associated nuclease, GFP = green fluorescent protein, URA = uracil selection marker. (B) Illustration of predicted colony outcomes for *S. cerevisiae* after receiving different plasmid combinations via conjugation. Control recipients (left) holding pSC5 and pV-GFP continue to grow on synthetic medium lacking uracil or on histidine-deficient medium supplemented with NTC. In contrast, recipients harboring pSC5 and pV-g1-7 (right) lose viability on yeast medium lacking uracil. Created with BioRender.com. pV-g1-7: CRISPR–Cas9 plasmids containing gRNAs. pV-GFP: CRISPR–Cas9 plasmid with no gRNA.

With this design, any loss of viability on –URA plates is linked to CRISPR activity rather than to defects in plasmid transfer. As has been shown in Figure 1B, control recipients carrying pSC5 and the non-targeting pV-GFP control are predicted to grow on –HIS/NTC and –URA medium, whereas recipients carrying pSC5 and a pV-g1-7 plasmids are predicted to lose viability on –URA medium but still grow on –HIS/NTC like the control. A similar idea was demonstrated by Cochrane et al. (2022), who used conjugation to deliver plasmids carrying restriction nuclease, into yeast, resulting in antifungal activity^21^. In our current work we aimed to have targeted elimination of yeast. To do that we designed seven guide RNAs targeting endogenous essential genes invoved in cell cycle control, ribosomal function, and DNA replication (Figure 2A). These included two guides targeting *Cell Division Cycle 28* (*CDC28*) (gRNA-1 and gRNA-2), which encodes a key regulator of the yeast cell cycle,^29^ and one guide each targeting *Ribosomal Protein S3* (*RPS3*) (gRNA-3), encoding a component of the small ribosomal subunit,^30^ *Ribosomal Protein S18B* (*RPS18B*) (gRNA-4), encoding another essential ribosomal protein,^31^ and *Minichromosome Maintenance Protein 2* (*MCM2*) (gRNA-7), which is required for DNA replication initiation.^32^ The remaining two guides (gRNA-5 and gRNA-6) were designed to target the intron of *Ribosomal Protein S4A* (*RPS4A*),^33^ enabling us to test whether cleavage of a non-coding region produces different response compared to targeting essential coding sequences (Figure 2B). These targets were selected because disruption of cell cycle regulation, DNA replication, or ribosome assembly is expected to strongly reduce viability, making them interesting candidates for antifungal killing. Among the coding-region guides, target positions are also different: gRNA-1, gRNA-2, and gRNA-7 cut near the start codon, while gRNA-3 and gRNA-4 target downstream regions (Figure 2B), enabling us to assess how cut-site position can impact lethality.

**Figure 2.**
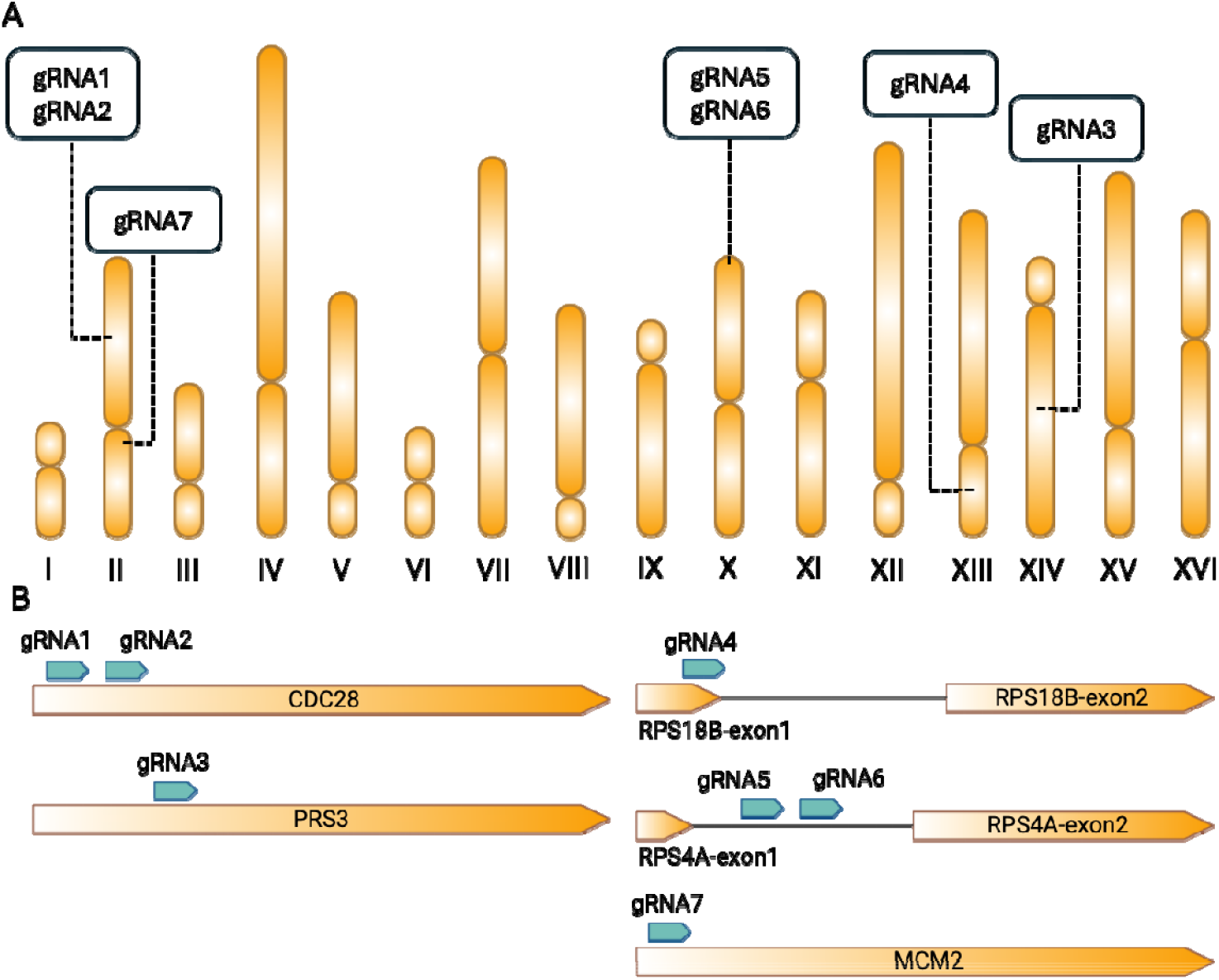
Genomic locations and gene-level positions of gRNA targets. (A) Schematic representation of the 16 *S. cerevisiae* chromosomes showing the approximate positions of all seven guide RNAs used in this study. (B) Gene-level maps showing the positions of individual gRNA binding sites relative to their corresponding coding regions or intron targets. Solid blocks represent exons, and connecting lines indicate introns. Arrows denote gRNA orientation. Gene diagrams are not drawn to scale.

Each guide was cloned into pV-GFP via Golden Gate assembly (pV-g1– pV-g7), and the resulting constructs were electroporated into *S. cerevisiae* to assess CRISPR-dependent growth defects (Figure 3). We used the pV-GFP plasmid (no gRNA) as a control to separate CRISPR-dependent lethality from any effects unrelated to the DNA cut (Figure 3). pV-g1 and pV-g7 showed the strongest lethality (∼97% and 100% reductions), whereas pV-g2 showed moderate suppression (∼66%) with noticeably smaller colonies, suggesting partial growth inhibition compared to the control. In contrast, pV-g3, pV-g4, pV-g5, and pV-g6 did not show any significantly different from the control. These results are consistent with previous reports that how guide position influences CRISPR efficiency. It has been shown that guides targeting different parts of the same gene can be different in their activity.^34^ Horlbeck *et al* (2016) also showed that local chromatin structure can limit Cas9 access at some sites, that reduces cleavage efficiency.^35^ Based on these results, we selected pV-g1, pV-g2, and pV-g7 for conjugation experiments and included pV-g6 (intron-targeting) as a non-lethal control.

**Figure 3.**
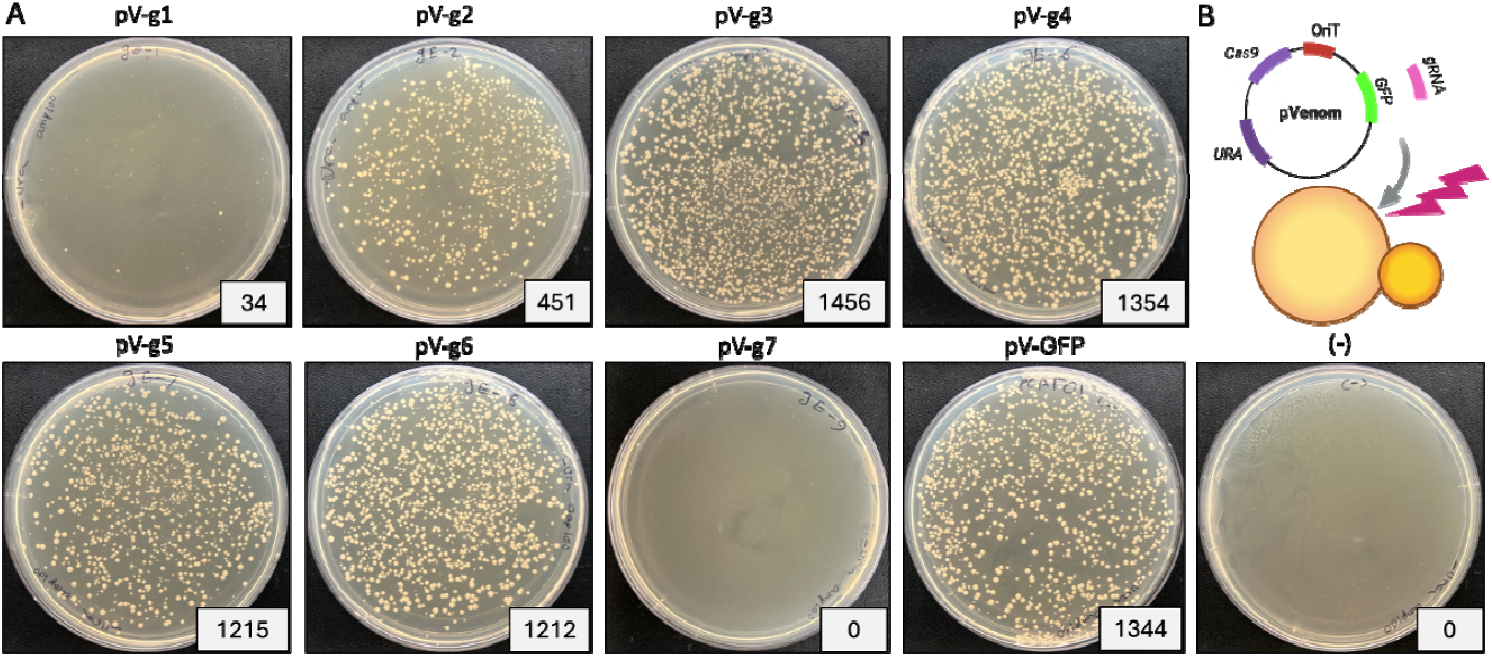
Evaluation of pVenom plasmids using electroporation as delivery into *S. cerevisiae*. (A) Yeast colony formation following transformation with the indicated pV-g1-7 or pV-GFP plasmids. Each plate shows the resulting colonies after selection on synthetic yeast media lacking uracil, and colony counts are indicated for each condition. (B) Schematic representation of yeast transformation with pVenom plasmids, illustrating plasmid entry and subsequent CRISPR–Cas9 activity. Note: The labels written directly on the plates during the experiment (numbered gE-1, 2, 5, 6, 7, 8, and 9) differ from the simplified plasmid names shown in the figure. For simplicity and consistency, the plate images were annotated using the standardized pV-GFP plasmid names (pV-g1, pV-g2, pV-g 3, pV-g 4, pV-g 5, pV-g6, and pV-g7).

We next evaluated conjugation method for delivering the pV plasmids to *S. cerevisiae* (Figure 4A,C). Plasmids pV-g1, pV-g2, and pV-g7 again produced strong colony formation suppression (87%, 99%, and 91%, respectively), while pV-g6, which targets an intron within *RPS4A*, produced no decrease which is consistent with the transformation results. Interestingly, gRNA-2 showed almost complete colony elimination (∼99%) when delivered by conjugation, while it showed only moderate lethality using delivery via electroporation. This difference is likely related to how the two delivery methods work and show difference in how many recipient cells successfully acquire and express the CRISPR plasmid. Recent work has also been shown that conjugation can successfully deliver CRISPR/Cas9 systems from bacteria into yeast, where they can efficiently cut episomal targets and suppress yeast growth. In these systems, Cas9 nucleases cut a plasmid carrying uracil selection marker, resulting in growth suppression on media lacking uracil^24^.

**Figure 4.**
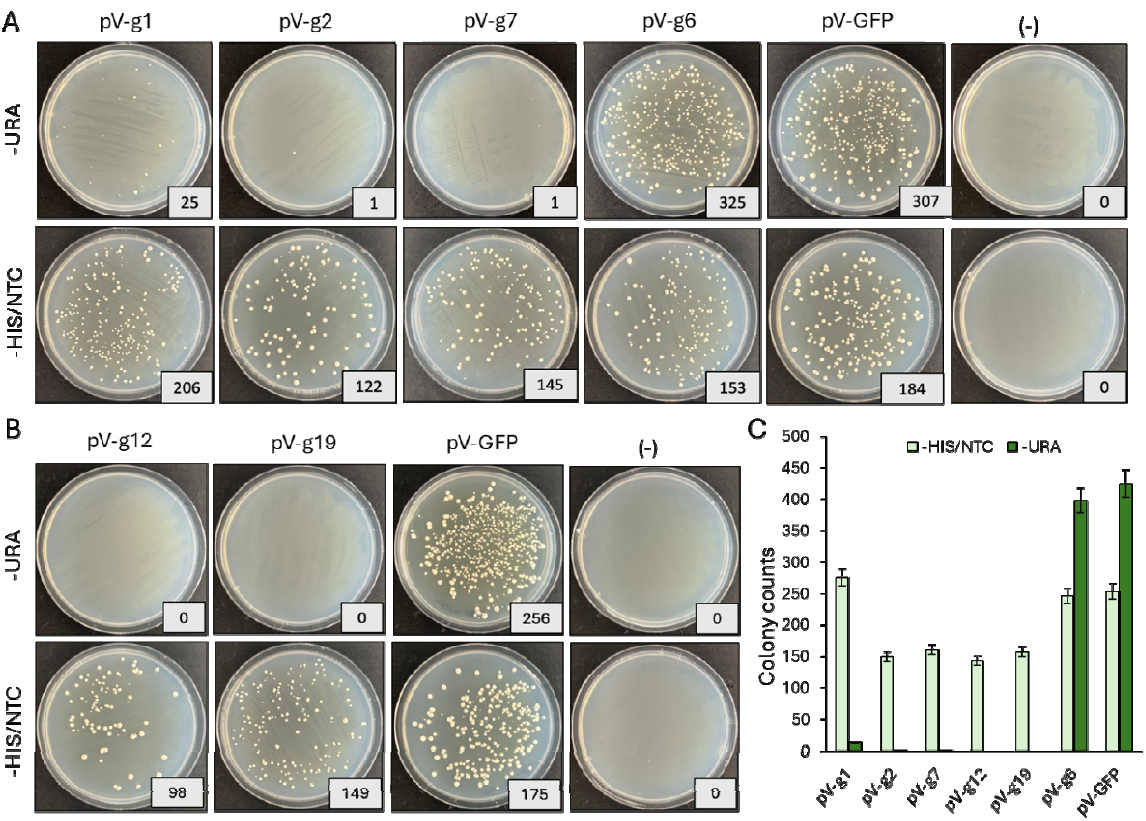
Conjugation-based evaluation of pVenom plasmids in *S. cerevisiae*. (A) Assessment of *S. cerevisiae* colony growth on –URA or –HIS/NTC plates after conjugative delivery of pVenom plasmids containing single-gRNA. (B) Assessment of *S. cerevisiae* colony growth on –URA or –HIS/NTC plates after conjugative pVe om plasmids with multiple gRNAs. (C) Colony counts of S. cerevisiae following conjugation with pVenom plasmids containing gRNA and the empty vector control. Bars indicate mean values from three independent experiments, and error bars show the standard error of the mean. -URA: synthetic yeast media lacking uracil; – HIS/NTC: synthetic yeast media lacking histidine supplemented with NTC (100 μg/mL).

Building on the single-guide results, we next constructed two dual-guide plasmids, pV-g12 (gRNA1 + gRNA2) and pV-g19 (gRNA1 + gRNA7), which were chosen based on their strong activity in initial conjugation assays. We then evaluated the multiplex constructs using conjugation, and interestingly, both constructs showed complete elimination of yeast colony formation compared to the control plasmid with no gRNA (Figure 4B,C). These results are in line with what has been reported previously., as Bao *et al*. (2014) and Jakočiūnas *et al*. 2015 showed that multiplex CRISPR editing improves penetrance and reduces escape events in *S. cerevisiae*.^36,37^ Taken together, these results suggest that multiple-guide pVenom plasmids are a stronger and more dependable antifungal approach than single-guide versions. Using multiple guides reduces the risk that one fails and increases the efficiency.

In conclusion, our results show that CRISPR plasmids delivered by conjugation can effectively kill *S. cerevisiae* when targeting endogenous genes, it also showed that guide activity depends on the essentiality of the target and where the cut is made. Furthermore, multiplex gRNA designs resulted in complete yeast colony elimination. Altogether, these findings demonstrate that COBRA technology should be further explore as a potential method for targeted fungal elimination.

## Method and materials

### Microbial strains and growth conditions

*Saccharomyces cerevisiae* strain VL6-48 (ATCC, catalog no. MYA-3666) was cultured in 2× yeast extract– peptone–dextrose (YPAD) medium supplemented with adenine hemisulfate (200 µg/mL; Sigma–Aldrich, #A2545, St. Louis, MO, USA) and ampicillin (100 µg/mL; BioBasic, Cat# AB0028, Canada). For selective growth, yeast was plated or cultured on one of the following media: (1) synthetic complete medium lacking histidine and supplemented with adenine hemisulfate (Takara, Inc., Cat# 630415, USA) and nourseothricin (100 µg/mL; Jena BioScience, Cat# AB-102XL, Germany).

*Escherichia coli* (Epi300, Epicenter) was grown at 37°C in Luria Broth (LB) supplemented with appropriate antibiotics (50 μg/ml Kanamycin and/or 40 μg/ml gentamycin). Cultures were incubated at 37°C, with liquid cultures shaking at 225 rotations per minute (rpm).

### Plasmid Construction

The pV-g1 plasmid was derived from the pWS158 plasmid^38^ by the addition of the *ACT1* yeast intron in the middle of the *Cas9* gene, the origin of transfer (oriT), and the *HIS3* gene using yeast assembly. pWS158 was amplified as two overlapping fragments. The origin of replication and the HIS gene were amplified as one fragment from pHflu3;^39^ and the *ACT1* intron was amplified from *S. cerevisiae* VL6-48 gDNA using primers that added 40 bp of overlapping homology to each end. (all primers are listed in Supplementary Table S1). The resulting amplicons were then combined in roughly equimolar amounts and co-transformed into *S. cerevisiae* spheroplasts, yielding the circular plasmid pV-GFP. Yeast assembly was conducted as previously described^40^ using mixtures of DNA fragments in place of a bacterial donor. Following assembly, DNA was isolated from pooled *S. cerevisiae* transformants and electroporated into *E. coli* Epi300. Individual colonies were screened using multiplex PCR.

### gRNA designing and Cloning

Guide RNAs (gRNAs) were designed to target essential genes involved in cell cycle regulation, ribosomal function, and DNA replication in *S. cerevisiae*. These targets included two gRNAs for *CDC28* (pV-g1 and pV-g2), and one each for *RPS3* (pV-g3), *RPS18B* (pV-g4), and *MCM2* (pV-g7), as well as two gRNAs targeting an intron within *RPS4A* (pV-g5 and pV-g6) (Supplementary Table S2). Oligonucleotides corresponding to the seven gRNA sequences were designed using Benchling,^38^ (Supplementary Table S3) phosphorylated with T4 Polynucleotide Kinase (PNK), and annealed. Briefly, each oligo (1 µL, 100 µM) was mixed with 1 µL 10× ligase buffer, 7 µL nuclease-free water, and 1 µL T4 PNK, and incubated for 1 h at 37°C. Equal volumes of the phosphorylated oligos were then combined and adjusted to a final volume of 200 µL with water. A 50 µL aliquot of this mixture was subjected to an annealing program consisting of 96°C for 6 min, a gradual ramp down at 0.1°C per second to 23°C, and a hold at 23°C. The annealed oligos were subsequently assembled into the pV-GFP vectors using Golden Gate assembly. Each 15 µL reaction contained 20 fmol of BsmBI-digested pV-GFP, annealed gRNA insert, 1 µL T7 DNA ligase (New England BioLabs, Cat. #M0202L, USA), and 1 µL BsmBI-HFv2 (New England BioLabs, Cat. #R3733S, USA). The reaction was cycled 10 times between 42°C for 2 min and 16°C for 5 min, followed by 60°C for 10 min, 80°C for 10 min, and a final hold at 12°C. The pV-GFP vector, which carries *Cas9, GFP, URA3*, and *HIS3*, was digested with BsmBI to remove *GFP* and replace it with each gRNA cassette. The resulting plasmids were transformed into *E. coli* EPI300, and non-fluorescent (non-GFP) colonies were selected on kanamycin-containing medium. Sequences of all assembled plasmids are provided in Supplementary Table S2.

### Synthesis and cloning of multiplex gRNAs

DNA fragments containing *gRNA1–gRNA2* and *gRNA1–gRNA7* cassettes were synthesized as gBlocks Gene Fragments (Integrated DNA Technologies, Coralville, IA, USA) (Table S2). The synthesized fragments were cloned into the pV-GFP vector as described in the previous Section. Sequences of all assembled plasmids are provided in Supplementary Table S2.

### Plasmid delivery via electroporation

Electroporation experiments were conducted as previously described.^41^ Briefly, a 50 mL yeast culture was grown in YPAD to an OD_600_ of 1.5 and pelleted at 5000 × g for 5 min. The pellet was resuspended in 10 mL of 0.1 M LiOAc in 1× TE (pH 7.5) and incubated at 30 °C with shaking at 150 rpm for 1 h. Subsequently, 250 µL of 1 M DTT was added, and incubation continued for 30 min. Cells were then washed sequentially with 40 mL and 25 mL of ice-cold sterile water, followed by 5 mL of ice-cold 1 M sorbitol, and finally resuspended in 250 µL of 1 M sorbitol. Electrocompetent cells were aliquoted (100 µL) and stored at −80 °C. For transformation, cells were thawed rapidly at 37 °C and kept on ice. Electroporation mixtures contained 40 µL of electrocompetent cells, 2 µL of plasmid DNA (400 ng), and 20 µg of single-stranded DNA in a 0.2 cm cuvette. Electroporation was performed at 1.8 kV, 200 Ω, and 25 µF. After pulsing, 1 mL of YPAD was added to each cuvette, and cells were transferred to Eppendorf tubes for recovery at 30 °C and 225 rpm for 2 h. Finally, 200 µL of each transformation was plated on selective media and incubated at 30 °C for 2 days.

### Conjugation

Conjugation happened in *trans-* orientation. *E. coli* harboring pSC5 and pV-g1-containing each gRNA. *E. coli* and *S. cerevisiae* strains were prepared in advance and stored as frozen aliquots for later use in conjugation assays. To prepare the *E. coli* cultures, overnight saturated cultures initiated from single colonies were diluted to an OD_600_ of 0.1 in 50 mL of LB medium containing the appropriate antibiotics (Table S2) and grown until reaching an OD_600_ of approximately 1.0. Cells were then harvested by centrifugation (3,000 × RCF, 15 min), resuspended in 500 µL of ice-cold 10% glycerol, and divided into 100 µL aliquots. The aliquots were stored at −80°C until needed. For the *S. cerevisiae* recipient strain, a single colony was inoculated into 5 mL of 2× YPDA medium supplemented with ampicillin (100 µg mL^−1^) and incubated for 7 h. This pre-culture was then diluted into 50 mL of fresh 2× YPDA medium with ampicillin (100 µg mL^−1^) and incubated until an OD_600_ of ∼3.0 was achieved (approximately 17 h). Cells were collected by centrifugation (3,000 × RCF, 5 min), resuspended in 500 µL of ice-cold 10% glycerol, aliquoted into 250 µL portions, and frozen in a −80°C ethanol bath, then stored at −80°C. On the day of conjugation, solid conjugation medium consisting of 20 mL per plate of 1.8% agar, 10% LB, and synthetic defined (SD) minimal glucose medium lacking histidine (Takara Bio, Cat. #630411, USA) was prepared and dried for 30 minutes. Frozen donor and recipient aliquots were thawed on ice for about 20 min. A 50 µL aliquot of *S. cerevisiae* cells was added to 100 µL of *E. coli* cells, mixed gently by pipetting, and the mixture was evenly spread on the conjugation plate for mating.

## Acknowledgments

This research was funded by Western Innovation Fund and Natural Sciences and Engineering Research Council of Canada (NSERC), grant number: RGPIN-2018-06172. In addition, this research was supported by Ontario Agri-Food Research Initiative (OAFRI - grant number OAF-2023-102672). OAFRI is supported by the Governments of Canada and Ontario through the Sustainable Canadian Agricultural Partnership (Sustainable CAP), a 5-year, federal-provincial-territorial initiative. Graphical abstract and figures 1-3 were partially created using Biorender.com.

## Author Contribution

VN: conceptualization, formal analysis, investigation, methodology, writing – original draft, writing – review & editing; BJK: conceptualization, formal analysis, funding acquisition, methodology, resources, supervision, writing – review & editing.

